# *PySilSub*: An open-source Python toolbox for implementing the method of silent substitution in vision and non-visual photoreception research

**DOI:** 10.1101/2023.03.30.533110

**Authors:** Joel T. Martin, Geoffrey M. Boynton, Daniel H. Baker, Alex R. Wade, Manuel Spitschan

## Abstract

The normal human retina contains several classes of photosensitive cell—rods for low-light vision, three cone classes for daylight vision, and intrinsically photosensitive retinal ganglion cells (ipRGCs) expressing melanopsin for non-image-forming functions including pupil control, melatonin suppression and circadian photoentrainment. The spectral sensitivities of the photoreceptors overlap significantly, which means that most lights will stimulate all photoreceptors, to varying degrees. The method of silent substitution is a powerful tool for stimulating individual photoreceptor classes selectively and has found much use in research and clinical settings. The main hardware requirement for silent substitution is a spectrally calibrated light stimulation system with at least as many primaries as there are photoreceptors under consideration. Device settings that will produce lights to selectively stimulate the photoreceptor(s) of interest can be found using a variety of analytic and algorithmic approaches. Here we present *PySilSub* (https://github.com/PySilentSubstitution/pysilsub), a novel Python package for silent substitution featuring flexible object-oriented support for individual colorimetric observer models (including human and mouse observers), multi-primary stimulation devices, and solving silent substitution problems with linear algebra and constrained numerical optimisation. The toolbox is registered with the Python Package Index and includes example data sets from various multi-primary systems. We hope that *PySilSub* will facilitate the application of silent substitution in research and clinical settings.

---

Human vision begins when light from a retinal image is absorbed by photopigments present in the outer segments of rod and cone photoreceptors in the retina (Grünert & Martin, 2020; Jindrová, 1998; Sung & Chuang, 2010). Quantal absorption within the photoreceptors gives rise to electrochemical signals whose integration in functionally and anatomically specialized pathways provides visual cortex with information on brightness, spatial frequency, colour, and other dimensions of visual experience (Hubel & Wiesel, 1962; Derrington et al., 1984; Krauskopf et al., 1982; Webster & Mollon, 1994). At scotopic light levels (e.g., starlight) vision is mediated primarily by rods. Though absent from the fovea, rods are most numerous of the photoreceptors and otherwise widely distributed across the retina. At photopic light levels (e.g., daylight), the rod photopigment becomes saturated, and it is the short- (S), medium- (M) and long- (L) wavelength light sensitive cones that mediate vision. Cone cells are packed densely into the fovea and distributed more sparsely elsewhere. At mesopic light levels (e.g., full moon, night-time urban lighting), vision is served by complex interactions of signals from rod and cone photoreceptors (Zele & Cao, 2015).

Humans also possess a third class of photoreceptor. A relatively small population of intrinsically photosensitive retinal ganglion cells (ipRGC) express the photopigment melanopsin in their axons and soma (Provencio et al., 2000). The ipRGCs provide an additional light-sensing pathway with an important role in “non-visual” functions, including circadian photoentrainment and control of pupil size, via their direct projections to the suprachiasmatic nucleus of the hypothalamus and the olivary pretectal nucleus of the midbrain, respectively (Gamlin et al., 2007; Ruby et al., 2002). Although ipRGCs do not mediate vision in the same way as rods and cones, laboratory studies have shown that melanopsin activation can facilitate the processing of cone signals and influence brightness perception (Brown et al., 2012; Allen et al., 2014; Davis et al., 2015).

As illustrated in Figure 1, the photoreceptor classes have different spectral sensitivities. The curve for each type of photoreceptor describes the probability of its capturing a photon at a given wavelength, so S cones are about 10 times more likely than L-cones to capture photons at 450 nm, and the likelihood of L and M cones capturing at 550 nm is roughly the same. Because the spectral sensitivities of the photoreceptors overlap, most lights in the visible spectrum will stimulate all types of photoreceptors, albeit to varying degrees. However, with the method of silent substitution, it is possible to prepare light stimuli that selectively target individual photoreceptor classes (Estévez & Spekreijse, 1982).

**Figure 1.**
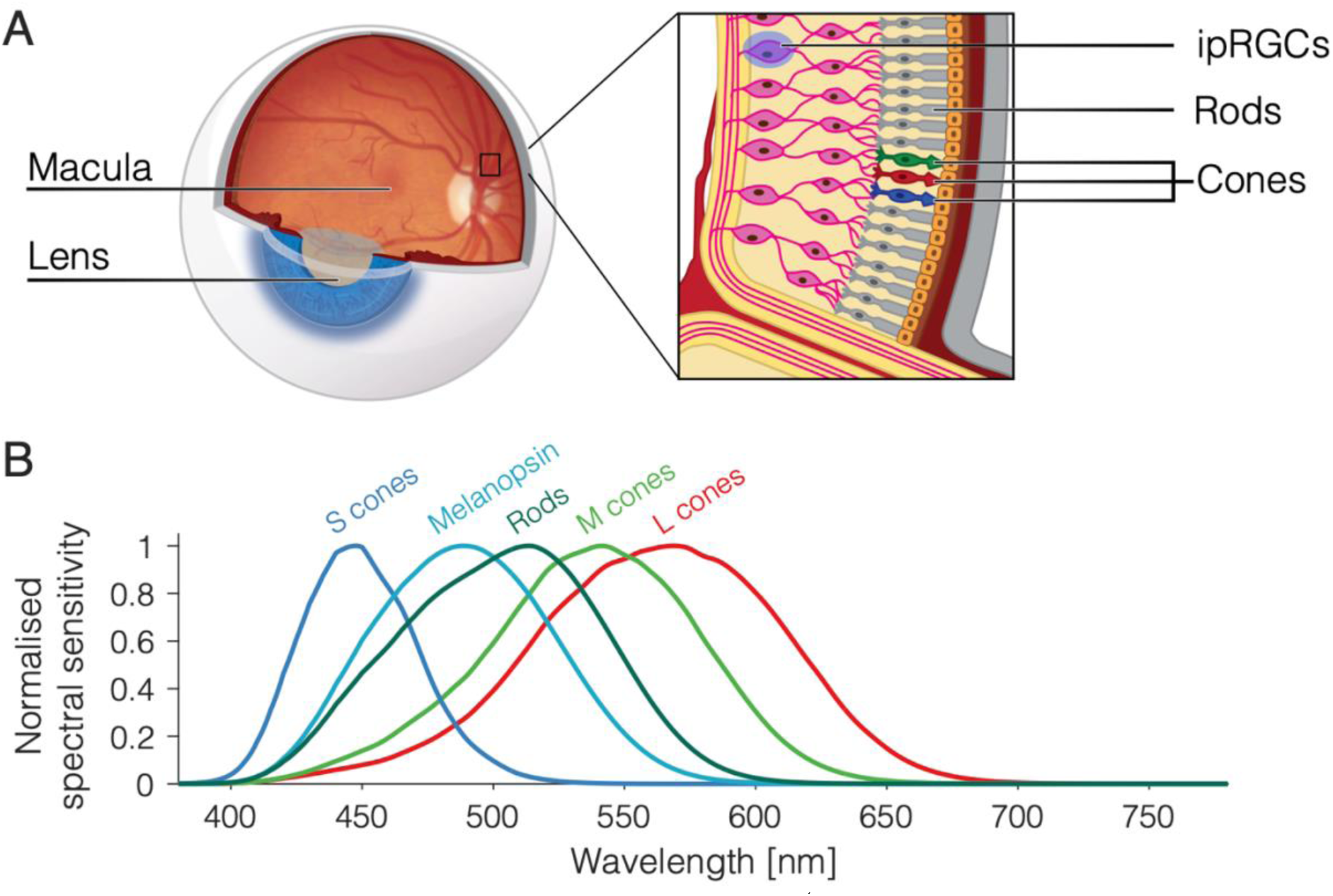
Photoreceptors and their spectral sensitivities (adapted from Figure 3 of Blume et al., 2019; reproduced under the CC BY 4.0 license, http://creativecommons.org/licenses/by/4.0/). (A) Human eye with inset showing the three types of retinal photoreceptor: rods, cones, and intrinsically photosensitive retinal ganglion cells (ipRGCs). (B) Overlapping spectral sensitivities of retinal photoreceptors. The curve for each photoreceptor describes the relative probability of photon capture at a given wavelength.

Silent substitution is an elegant technique that involves using a spectrally calibrated multiprimary stimulation system to produce spectra that selectively stimulate one class of retinal photoreceptor whilst maintaining a constant level of activation in the others. The method exploits Rushton’s (1972) principle of univariance, which states that the output of a photoreceptor is one-dimensional and depends upon quantum catch, not upon which quanta are caught. In other words, different spectra will have an identical effect on a photoreceptor providing they lead to the same number of photons being absorbed. The principle of univariance and its relevance to silent substitution is covered in greater detail by Estévez and Spekreijse (1982), along with a treatment of the method’s history. In vision science, silent substitution has contributed to our understanding of human colour vision mechanisms (Horiguchi et al., 2013) and it has enabled researchers to examine how targeted photoreceptor stimulation affects physiological responses such as the electroretinogram (Maguire et al., 2017), and the pupillary light reflex (Spitschan et al., 2014).

The method of silent substitution can be used to generate metamers: spectra that have different wavelength distributions, but which produce the same cone excitation (the principle underlying all modern colour display devices). Such lights can be used to stimulate pathways contributing to “non-visual” responses to light, including melatonin suppression (Allen et al., 2018; Blume et al., 2022; Souman et al., 2018), sleep (Blume et al., 2022; Schöllhorn et al., 2023), and other neuroendocrine and circadian functions (Zandi et al., 2021). The method of silent substitution can also be used to investigate the contribution of melanopsin signaling to canonical visual processing (Allen et al., 2019; Brown et al., 2012; DeLawyer et al., 2020; Spitschan et al., 2017; Uprety et al., 2022; Vincent et al., 2021), and its potential as a diagnostic tool for retinal disease has garnered attention in recent years (Kuze et al., 2017; Wise et al., 2021).

The main hardware requirement for silent substitution is a spectrally calibrated stimulation system with at least as many primaries as there are photoreceptors of interest. With three cone classes in addition to rods and ipRGCs, this generally means that five primaries are needed, but four may suffice when working in the photopic range and sufficient measures are taken to ensure the rods are saturated (see Adelson, 1982; Aguilar & Stiles, 1954; Kremers et al., 2009; Shapiro, 2002; Sharpe et al., 1989). The primaries should be independently addressable, additive, and ideally stable over time with a linear input-output function. Peak wavelength and bandwidth of the primaries are key considerations that will define the gamut and available contrast (Evéquoz et al., 2021), and the light source should be integrated into an optical system for stimulus delivery (Barrionuevo et al, 2022)—usually a Ganzfeld (e.g., Martin et al., 2021; Geiser et al., 2019), specialised Maxwellian view (e.g., Cao et al., 2015; Nugent & Zele, 2022), projector (Allen et al., 2018; Allen et al., 2019; DeLawyer et al., 2020; Hexley et al., 2020; Spitschan et al., 2019; Yamakawa et al., 2019) or display (Blume et al., 2022; Schöllhorn et al., 2023) system. In a review on stimulation devices used in the literature, Conus and Geiser (2020) found that in most cases the device had 4 or 5 primaries and was built from scratch using LEDs, optical bench components, and microprocessors (e.g., Arduino) with intensity controlled by pulse width modulation.

Also required for silent substitution are estimates of the photoreceptor action spectra of the observer, which, in humans, vary even among colour-normal observers due to differences in ocular physiology. Before striking the retina, incident light must first travel through the lens and ocular media pigment of the observer, which act as prereceptoral filters to effectively shift the spectral sensitivity of the underlying photoreceptors. The crystalline lens of the eye accumulates its yellow pigment with age due to absorption of near-UV radiation (Norren & Vos, 1974; Pokorny et al., 1987; Weale, 1988) and a yellower lens transmits less short-wavelength light. The macula pigment is a yellow carotenoid spot that sits above foveal photoreceptors and reduces the spectral sensitivity of underlying cones in a manner that becomes more pronounced for smaller stimulus field sizes (Chen et al., 2001; Whitehead et al., 2006). Observer age and stimulus field size therefore combine to alter the effective spectral sensitivity functions of a given observer under particular viewing conditions (Figure 2; see also Figure 5, A-B).

**Figure 2.**
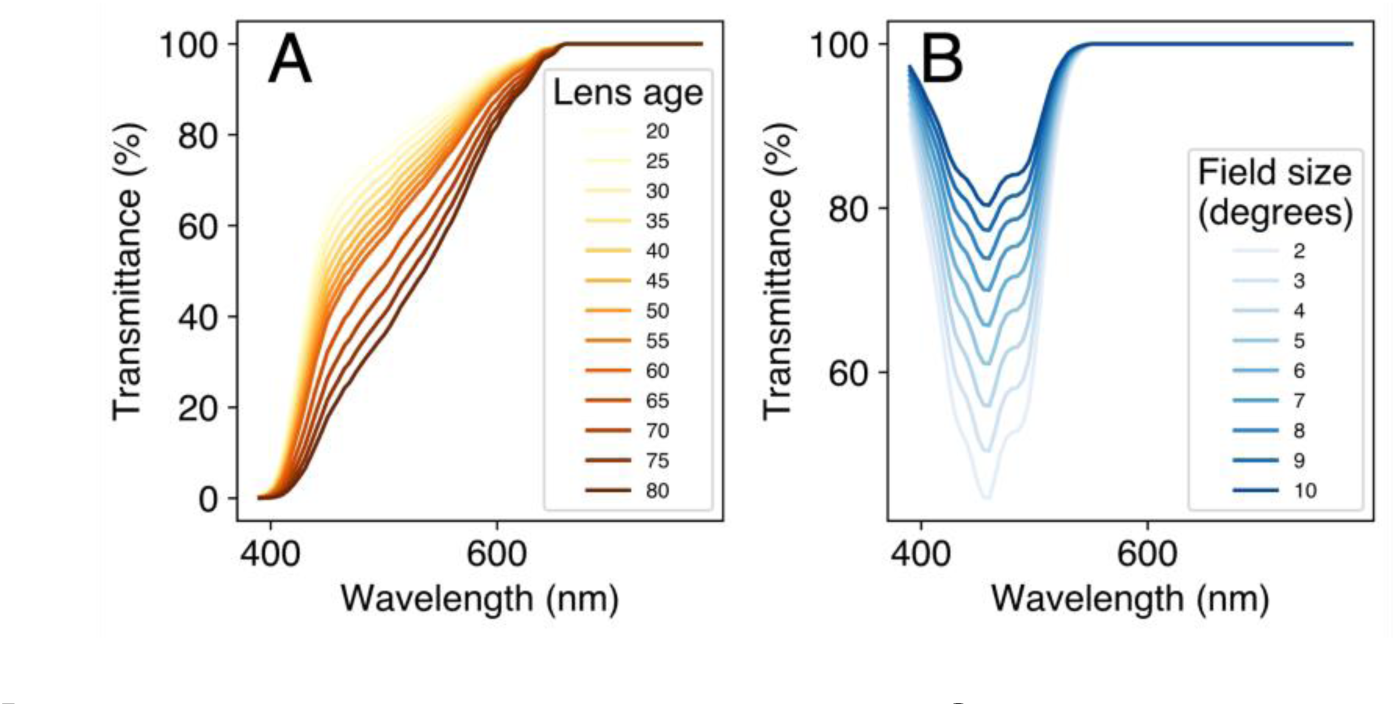
Prereceptoral filtering of light by ocular media. Spectral transmittance functions of (A) the human lens for ages 20 to 80 years (older lenses transmit less light), and (B) the macula pigment for field sizes 2° to 10°. The human lens naturally accumulates a yellow pigment with age due to absorption of near-UV radiation (Lerman, 1980; Feldman et al., 2022) and the macula pigment is a yellow carotenoid spot at the position of the fovea (see Grünert & Martin, 2020, Figure 1). All transmittance functions were calculated in accordance with CIEPO06 (CIE, 2006).

The International Commission on Illumination (CIE) defines average colorimetric observer models with estimates of photoreceptor spectral sensitivities for a given age and field size. The CIE observers are based on decades of research involving predominantly psychophysical methods (Smith & Pokorny, 1975; Stiles & Burch, 1959; Stockman et al., 1993, 1999; Stockman & Sharpe, 2000; Vos & Walraven, 1970) but also techniques such as suction electrode recording and microspectrophotometry of photoreceptors (e.g., Baylor et al., 1984; Bowmaker et al., 1978; Bowmaker & Dartnall, 1980). The CIE 2006 Physiological Observer (developed in CIE 170-1:2006 and abbreviated here as CIEPO06: CIE, 2006) defines fundamental cone spectral sensitivity functions for 2° and 10° field sizes and outlines a framework for calculating average spectral sensitivity functions for any age between 20 and 80 years and any field size between 1° and 10°. More recently, prompted by the pioneering research into melanopsin-containing ipRGCs (e.g., Al Enezi et al., 2011; Berson et al., 2002; Brown et al., 2013; Dacey et al., 2005; Gamlin et al., 2007; Lucas et al., 2014; Provencio et al., 2000; Spitschan, 2019), the CIE released an International Standard—CIE S 026/E:2018 (CIE, 2018)—that includes the melanopic and rhodopic action spectra alongside the CIEPO06 cone action spectra and outlines best practices for documenting photoreceptor responses to light.

With an accurately calibrated multiprimary system, one can predict *E*(*λ*), the spectral output of the device (radiance or irradiance, in energy units), for any combination of primary gain settings using interpolation followed by a linear combination of primary spectra. Then, with photoreceptor action spectra appropriate to the intended observer, one can compute the activation level for each class of photoreceptors following the equation:

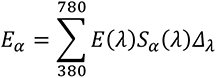

*Equation 1*. Calculating α-opic irradiance

where *E_α_* is a scalar representing the photoreceptor activation for photoreceptor class α (e.g., one of the five photoreceptors: S cones, M cones, L cones, rods, ipRGCs), *S_α_*(*λ*) is the estimated photoreceptor action spectrum, and *Δ_λ_* is the width of the spectral sampling. By convention (CIE, 2018; Lucas et al., 2014), when this computation concerns the spectral sensitivity functions of retinal photoreceptors, the resulting measures may be called α-opic irradiance. Figure 3 shows an example spectral measurement alongside the α-opic action spectra and the resulting α-opic irradiance measures that follow from Equation 1.

**Figure 3.**
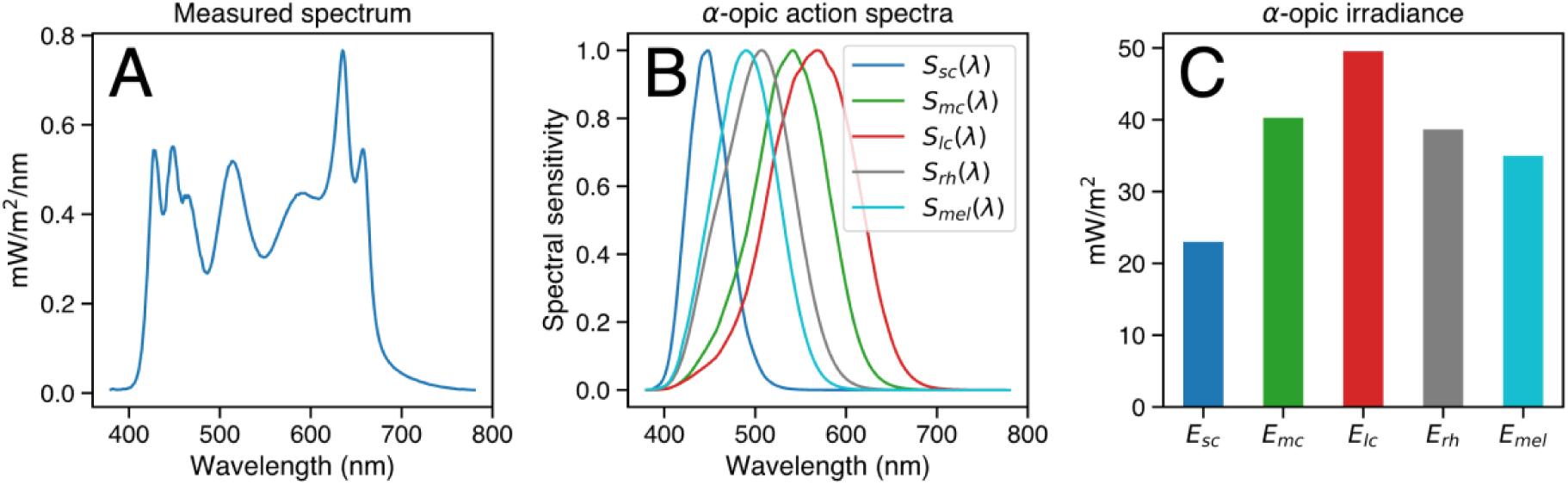
Calculating α-opic irradiance in accordance with CIE S 026 (CIE, 2018). (A) A spectral measurement is weighted against each of the (B) photoreceptor sensitivity functions to obtain (C) the respective α-opic quantities, following Equation 1.

The goal of silent substitution is to invert this process to determine the device input settings to produce two spectra—a background spectrum and a modulation spectrum— where the resulting α-opic quantities will be the same for all but the targeted photoreceptor(s). In the case where linearity holds across the entire α-opic computation, silent substitution is typically computed using linear algebra, by inverting a fully- or over-determined set of equations describing the mapping from primaries to photoreceptor responses (e.g., Cao et al., 2015; Maguire et al., 2016; Shapiro et al., 1996). However, where linearity fails at some point in the system—for example, if the spectrum of a particular primary changes as a function of its intensity—then it may still be possible to obtain a solution using constrained numerical optimisation (e.g., Spitschan et al., 2015).

In this Methods paper we introduce *PySilSub* (Martin et al., 2023), a Python toolbox that eases the computational burden of silent substitution by taking a structured approach to dealing with the various confounds and challenges outlined above. The toolbox provides flexible object-oriented support for multiprimary stimulation devices and individual colorimetric observer models, example calibration data for a range of devices, an intuitive interface for defining and solving silent substitution problems with linear algebra and constrained numerical optimization, and methods for accessing relevant standards and visualising solutions. The code is actively maintained on GitHub (https://github.com/PySilentSubstitution/pysilsub) under the MIT License and comes with a comprehensive testing suite and extensive online documentation with detailed examples. It is also registered with the Python Package Index (https://pypi.org/project/pysilsub/) and can therefore be installed with the *pip* packaging tool. Here, we provide an overview of the toolbox, describe key features, and demonstrate how it can be used.

## Overview

*PySilSub* (Martin et al., 2023) is written in *Python3* and depends on widely used libraries from the scientific Python ecosystem (*numpy*: Harris et al., 2020; *matplotlib*: Hunter, 2007; *pandas*: McKinney, 2010; *scipy*: Virtanen et al., 2020), and optionally some functionality from the *colour-science* package (Mansencal et al., 2022). Three core modules support colorimetric observer models (observers.py), multiprimary stimulation devices (devices.py), and defining, solving and visualising silent substitution problems (problems.py). Additional modules support CIE standards (CIE.py), prereceptoral filters (preceps.py), color computations (colorfuncs.py) and stimulus design (waves.py). Figure 4 lays out the anatomy of the package and highlights key features. The following sections describe the rationale and implementation of the core modules and key features.

**Figure 4.**
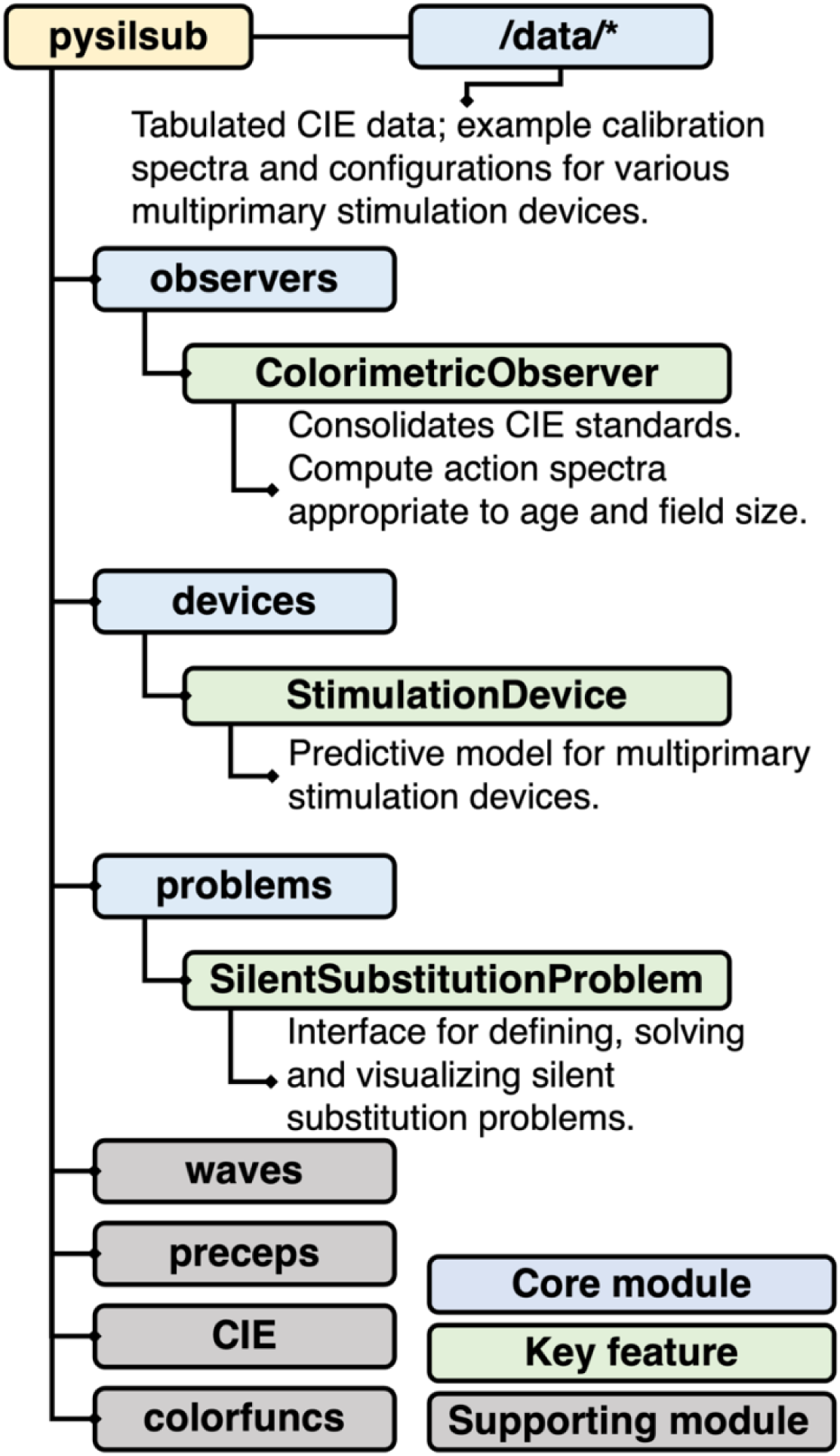
Anatomy and key features of *PySilSub*.

### Colorimetric observers (*observers.py*)

*PySilSub* consolidates the CIEOP06 (CIE, 2006) and CIE S 026 (CIE, 2018) observers in a *ColorimetricObserver* class that internally computes photoreceptor action spectra appropriate to a given age and field size. LMS cone fundamentals are constructed from the photopigment absorbance spectra (Stockman & Sharpe, 2000), taking account of the peak axial density of the respective photopigments, as well as lens (Pokorny et al., 1987; Stockman et al., 1999; Stockman & Sharpe, 2000) and macular pigment (Bone et al., 1988; Stockman et al., 1999) density, in accordance with the CIEPO06 model (CIE, 2006). The melanopic and rhodopic action spectra of the 32-year-old standard observer are taken from CIE S 026 (CIE, 2018) and adjusted for lens transmittance with a spectral correction function, in accordance with the standard. Note that, for consistency with the melanopic existing standards, a slightly different lens density function is used by default to correct the melanopic and rhodopic action spectra (for more information, see CIE S 026, CIE, 2018). Macular pigment correction is not applied to the rhodopic and melanopic action spectra as rods are not present at the fovea and ipRGCs are positioned above the retinal pigment layer (Trieschmann et al., 2008).

The *ColorimetricObserver* class stores the photoreceptor action spectra in a *pandas DataFrame* (McKinney, 2010) which interacts with other parts of the toolbox as required. This enables the toolbox to flexibly adapt to scenarios that may require alternative action spectra, such as targeting cones in the shadow of retinal blood vessels (Figure 5, C: Adams & Horton, 2002; Spitschan et al., 2015), performing silent substitution with non-human animals (Figure 5, D: e.g., Allen & Lucas, 2016; Mouland et al., 2019, 2021; Wise et al., 2021), or using alternative action spectra for any conceivable reason (e.g., to account for individual differences in spectral sensitivity arising from opsin gene polymorphisms), in which case one need only create a modified observer class with the relevant photoreceptor action spectra (see examples online). Incidentally, the *ColorimetricObserver* class is also a convenient open-source means of obtaining a full set of tabulated CIEPO06- and CIE S 026-compliant photoreceptor action spectra for a human observer for any age between 20 and 80 years and for any field size from 1° to 10°. Note that for field sizes larger than 10° the effects of the macula pigment are considered negligible, so one can safely assume a 10° field size.

**Figure 5.**
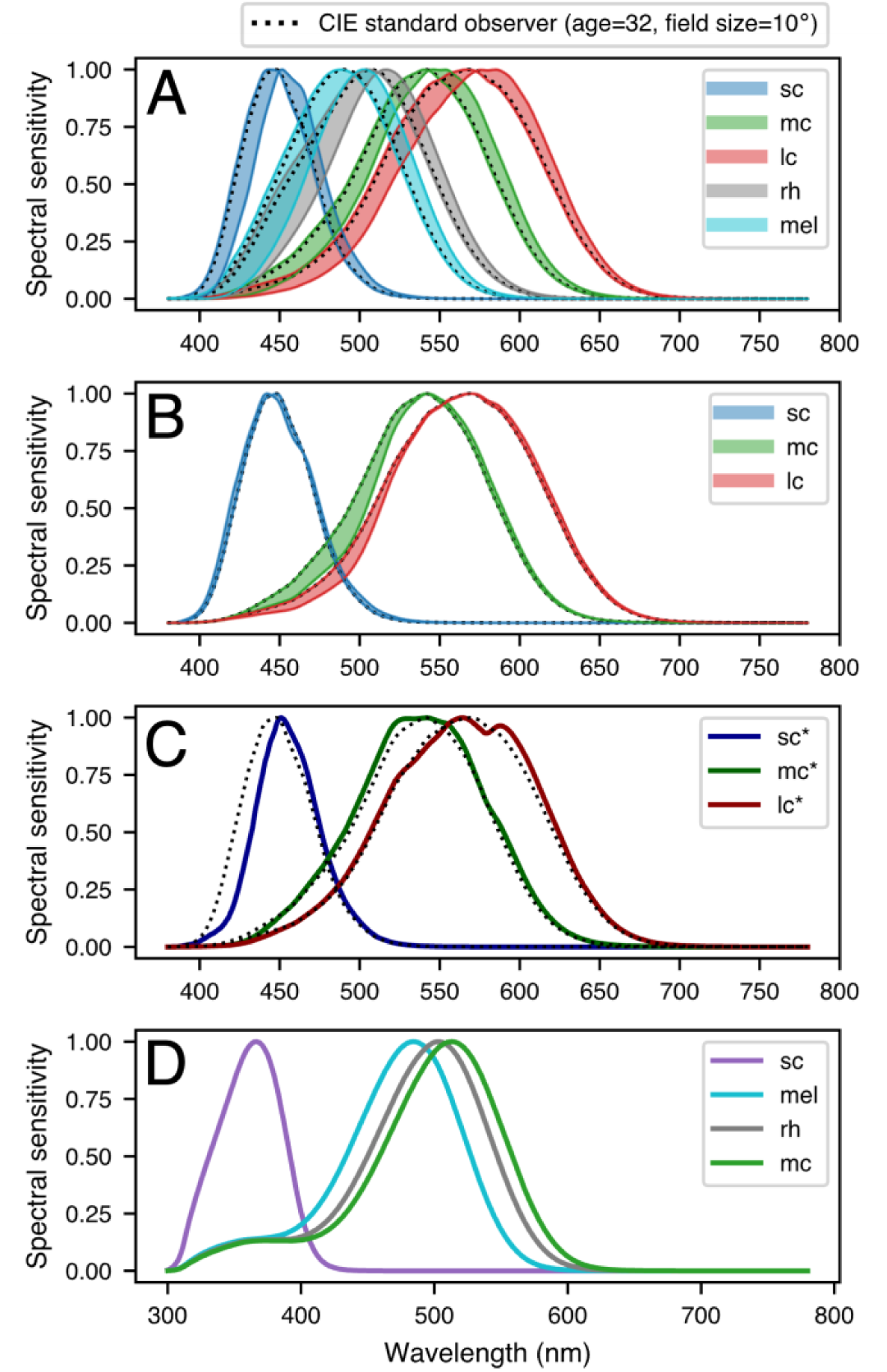
Spectral sensitivity of photoreceptors depends on the ocular physiology of the observer. (A) Lens yellowing with age causes a rightward shift in peak sensitivity of photopigments and alters the overall shape of sensitivity functions (shaded areas show the range of the spectral shift for ages 20 to 80 years). (B) Decreasing stimulus field size causes a small shift in S cone sensitivity and a marked dampening of the left-hand slopes of the sensitivity functions for M and L cones (shaded areas show the range of the spectral shift for field sizes 10° to 2°). Note that the effects of macular pigment absorption are considered negligible for rhodopsin and melanopsin, as rods are absent from the fovea and ipRGCs sit above the retinal pigment layer (Trieschmann et al., 2008). (C) Cones in the shadow of retinal blood vessels (sc*, mc*, lc*) have altered spectral sensitivity functions due to prefiltering of light by blood. Targeting these cones produces the entopic Purkinje tree percept (Adams and Horton, 2002; Spitschan et al., 2015). (D) Spectral sensitivity functions are species dependent. The example shown is for mice (from the Rodent Toolbox v2.2: Lucas et al., 2014). Note that mice do not possess a third cone and that the spectral sensitivity of photoreceptors peaks at different locations in the visual spectrum in comparison to humans. S cone sensitivity is shifted leftwards towards the near-UV portion of the spectrum and there is increased overlap of the M cone, melanopic and rhodopic action spectra.

### Multiprimary stimulation devices (*devices.py*)

*PySilSub* supports multiprimary stimulation devices through a generic *StimulationDevice* class that, given a set of accurate spectral calibration measurements, serves as a forward model for the device. The required format of the calibration data is a CSV file where the first row describes the wavelength sampling (e.g., 380, 381, … 780) and every other row is a spectral measurement (see Table 1) with units of radiance (e.g., *W*·*m*^−2^·*sr*) or irradiance (e.g., *W*·*m*^−2^·*nm*) in energy units. Note that if measurements are available in photon units only, they should be converted to energy units for agreement with the photoreceptor action spectra. Also required in the first row of the CSV file are the column headers *Primary* and *Setting*, whose corresponding values identify the source of the spectral measurements. As devices may not be linear it is prudent to obtain measurements across the entire range of primary intensities. If sampling with an OceanOptics (Ocean Insight Inc., Oxford, UK) or JETI (JETI Technische Instrumente, GmbH, Jena, Germany) spectrometer, one may be able to automate these measurements with the help of interfaces in Python packages such as *LuxPy* (Smet, 2020), *Seabreeze* (Poehlmann, 2019) and our own *PyPlr* (Martin et al., 2021; Martin & Spitschan, 2021).

**Table 1.**
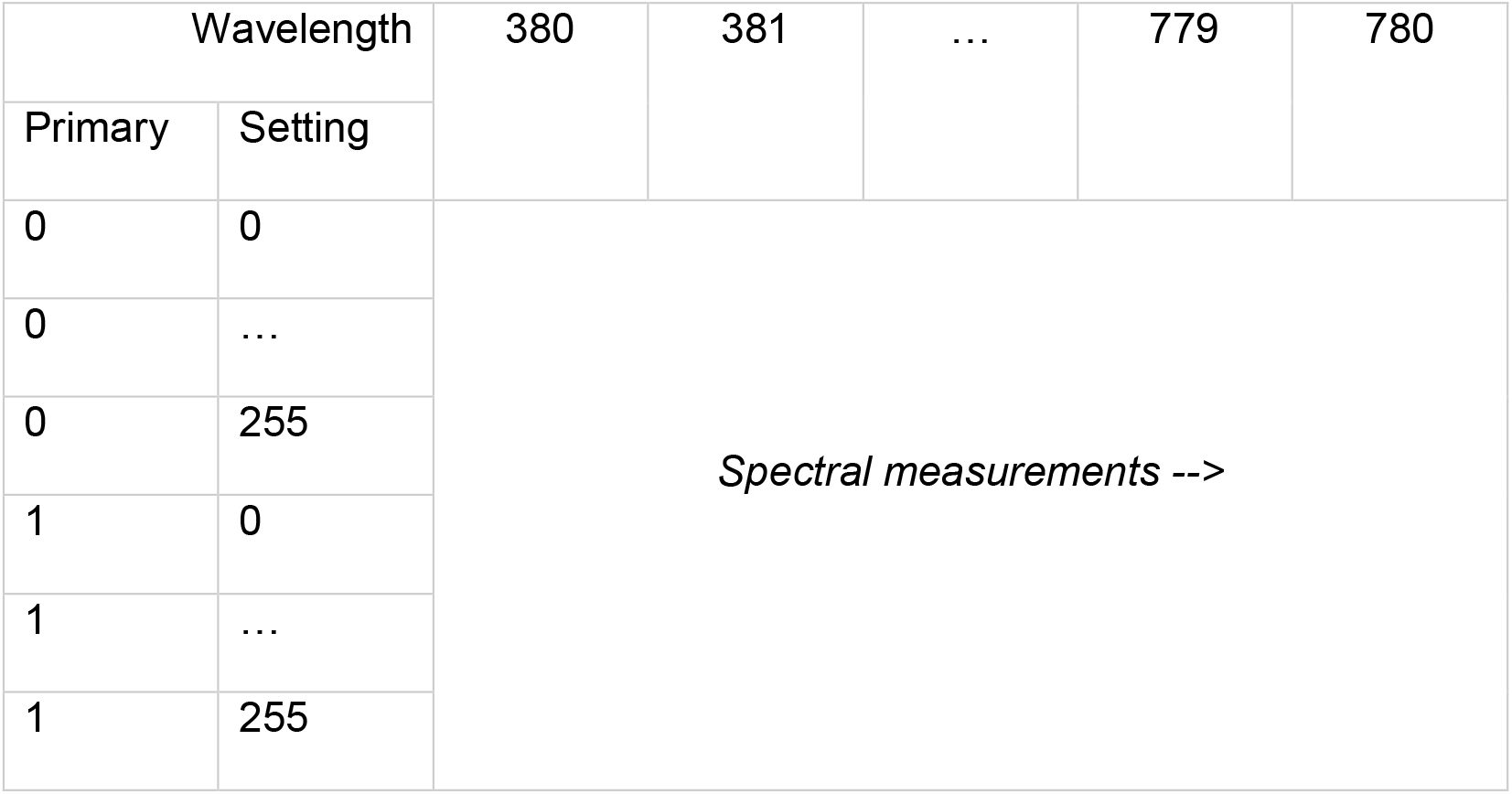
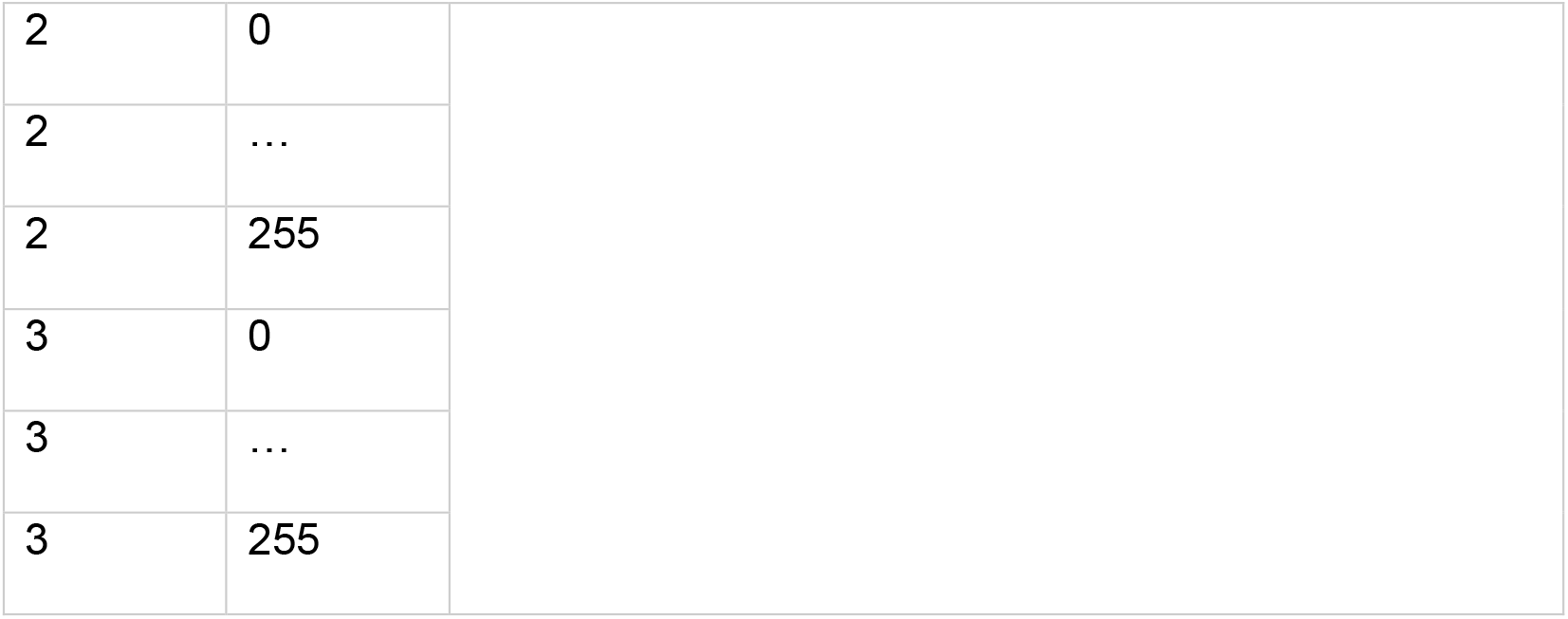
Calibration file format. A minimal example is shown for a hypothetical 4-primary device with 8-bit resolution. Each primary is assigned an ordinal index starting at zero and has a spectral measurement for the maximum (255) and minimum (0) input setting, where the minimum setting should reflect the ambient spectral power distribution (i.e., when all channels are off). A reasonable number of intermediary measurements should also be included to account for non-linearity in the device.

*StimulationDevice* has methods for predicting and visualizing the spectral and colorimetric output of a device, each of which responds to native device settings (e.g., 0-255 for a device with 8-bit resolution) or floating-point input weights between 0.0 and 1.0. The basis of all forward-predictions is a method called *.predict_primary_spd(…)* which uses input-wise linear interpolation on the calibration spectra to predict the spectral power distribution for a given primary at a given setting. A similar method called *.predict_multiprimary_spd(…)* returns the spectral power distribution for a linear combination of all primaries. Following the same pattern, *.predict_multiprimary_aopic(…)* computes a mutiprimary spectrum and returns the predicted α-opic irradiances for the assigned observer. By default, this is the CIEPO06 Standard Observer (age=32, field size=10°), but a custom observer may be assigned when building the class. Additional methods make it easy to perform gamma correction, change back and forth between weights and settings, visualize the calibration SPDs and gamut of the device, and instantiate from a preconfigured JSON file. Note that *PySilSub* includes example data for a range of multiprimary systems (Figure 6), which makes it easy to explore how the toolbox functions without having to develop one’s own system.

**Figure 6.**
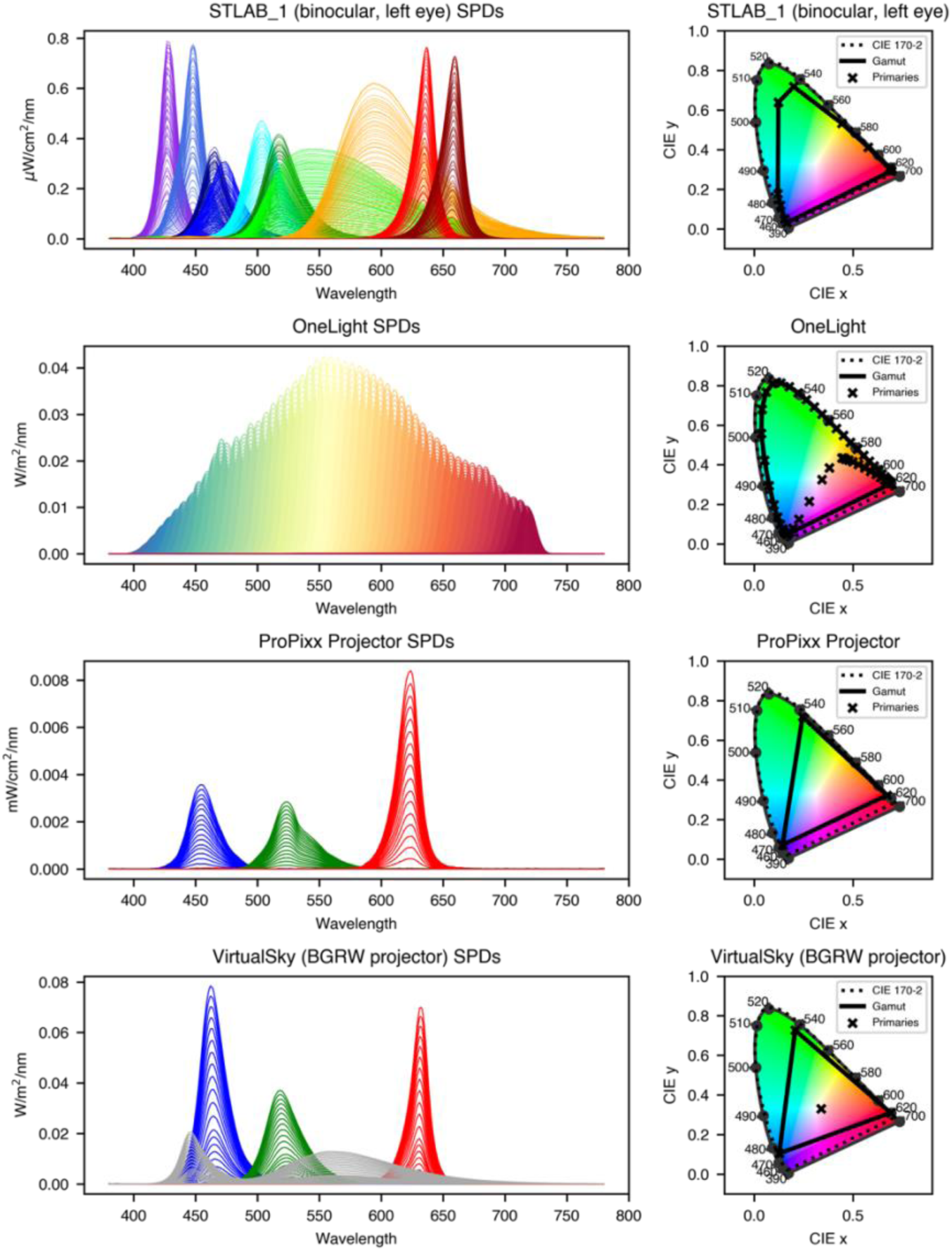
Example package data for multiprimary systems: (A) SpectraTuneLab 10-primary light engine (LEDMOTIVE technologies, LLC, Barcelona, Spain), (B) VISX Spectra Digital Light Engine (OneLight Corporation, Vancouver, Canada; calibration spectra from Spitschan et al., 2015), (C) VPixx ProPixx projector (VPixx Technologies Inc., Quebec, Canada), (D) BGRW VirtualSky (Fraunhofer IAO, Stuttgart, Germany; Stefani et al., 2012).

### Silent substitution problems (*problems.py*)

Stimuli in a silent substitution protocol typically take the form of pulses (e.g., Maguire et al., 2017; McAdams et al., 2018) or sinusoidal modulations (e.g., Barrionuevo et al., 2018; Spitschan et al., 2014) of photoreceptor-directed contrast presented against a background spectrum to which an observer has adapted. The background spectrum serves to maintain a set pattern of photoreceptor activations and the modulation spectrum increases activation of the targeted photoreceptor(s) without altering activation of the others. Michelson contrast, 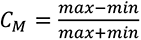, is the preferred metric for describing photoreceptor activations in bipolar modulations around a mean background, whereas Weber contrast, 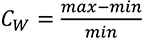 is more suitable for describing unipolar pulses against a relatively dim background. For transparency and consistency in silent substitution studies, it is important to state which contrast definition is used (Conus & Geiser, 2020).

Finding the native device inputs for the background and modulation spectra is a computational problem whose definition depends on one’s research aims and stimulus requirements. Above all, one must nominate which photoreceptor(s) to target, which to silence, and which to ignore (if any, see discussion), but it may also be advantageous to refine the problem by specifying a particular background spectrum, how much and of what kind of contrast to aim for on the targeted photoreceptor(s), and whether to impose any bounds on the primary inputs. These decisions then guide the development of a computational algorithm whose execution will return a solution—a set of native device settings for the background and modulation spectra—that satisfies the constraints in the problem specification.

With *PySilSub*, this process is streamlined by a *SilentSubstitutionProblem* class that inherits from *StimulationDevice*, has a *ColorimetricObserver* assigned to it, and incorporates properties for defining, and methods for solving and visualizing, silent substitution problems. After instantiating, a problem is conditioned by setting several basic and well-documented properties (i.e., managed attributes), including which photoreceptors to target, silence and ignore, and optionally the background spectrum, target contrast and primary input bounds. A solution is then obtained with a call to one of the two solver methods described below (see Figure 7 for a set of examples).

**Figure 7.**
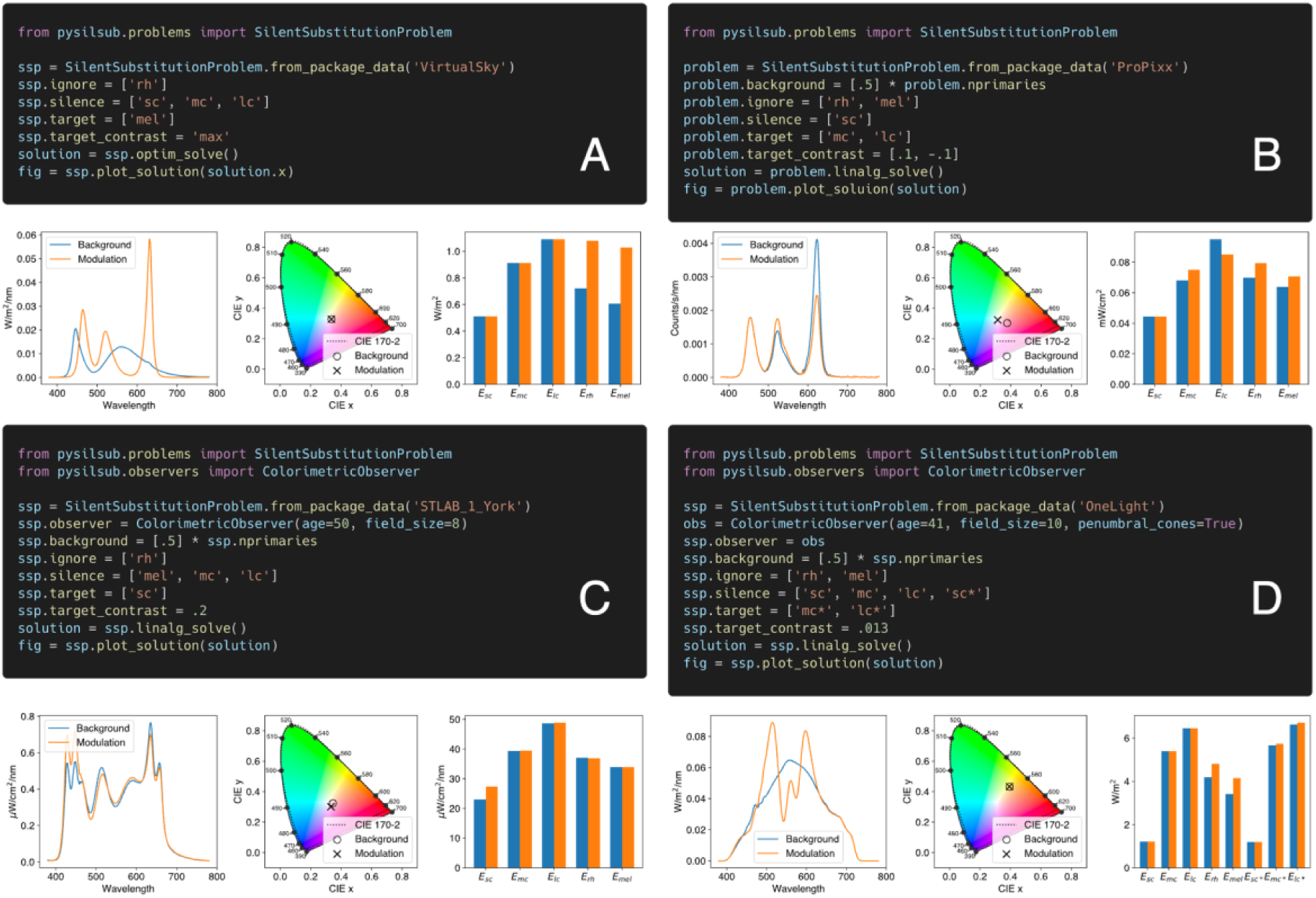
Defining and solving silent substitution problems with *PySilSub*’s example package data. (A) Using a BGRW VirtualSky (Fraunhofer IAO, Stuttgart, Germany; Stefani et al., 2012) to maximize melanopic contrast whilst ignoring rods and minimizing cones. Solved with numerical optimization without having specified a background spectrum (the background is optimized and included in the solution). (B) Using a VPixx ProPixx projector to target the L-M postreceptoral pathway whilst minimizing S cone contrast and ignoring rods and melanopsin, against a background spectrum defined as each of the three primaries at their half-maximum setting. Solved with linear algebra. (C) Using a SpectraTuneLAB 10-primary light engine (LEDMOTIVE technologies, LLC, Barcelona, Spain) to target S cones with 20% contrast, for a 50-year-old observer and field size of 8 degrees, whilst ignoring rods and minimizing melanopsin and M/L cones, against a background spectrum defined as all primaries at their half-maximum setting. Solved with linear algebra. (D) Using a VISX Spectra Digital Light Engine (OneLight Corporation, Vancouver, Canada; calibration spectra from Spitschan et al., 2015) to target M and L cones in the shadow of retinal blood vessels for a 41-year-old observer and a field size of 10 degrees.

### Linear algebra: .linalg_solve()

Linear algebra is a fast and efficient means of solving silent substitution problems (Cao et al., 2015; Evéquoz et al., 2021; Shapiro et al., 1996). The *.linalg_solve()* method applies specifically for situations where the background spectrum and target contrast are already known. To illustrate how it works, suppose we have a 5-primary system and that for a background spectrum we wish to use the mixture of all primaries at half-power, *α_bg_* = [.5 .5 .5 .5 .5]. First, we generate the matrix *P_bg_*, where the rows of *P_bg_* are the predicted spectral power distributions for each primary component of the background spectrum P_bg_ = [*p*_1_(*λ*) *p*_2_(*λ*) *p*_3_(*λ*) *p*_4_(*λ*) *p*_5_(*λ*)]. We then compute the matrix of *α*-opic irradiances:

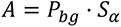

where the columns of *S_α_* are the photoreceptor action spectra of the observer, *S_α_* = [*sc*(*λ*) *mc*(*λ*) *lc*(*λ*) *rh*(*λ*) *mel*(*λ*)]*^T^*. Given a set of scaling coefficients for the primaries, *α*_sc_ = [*p*_1_ *p*_2_ *p*_3_ *p*_4_ *p*_5_], the forward model predicts that the output modulation *β* = [*sc mc lc rh mel*] is *β* = *α*_sc_*A*. To invert this process, we calculate *α_sc_* = *βA*^−1^, which provides the desired scaling coefficients that must be added to the primary weights for the background to produce the desired output modulation *β*. Finally, the solution is given as, *α_mod =_ α_bg_ = α_sc_*.

The *.linalg_solve()* method relies on functionality from *numpy*’s *linalg* module (Harris et al., 2020). Specifically, in the case of square matrices (e.g., five primaries and five photoreceptors) the *inv* function is used to perform matrix inversion, whereas non-square (i.e., overdetermined) matrices require the *pinv* (Moore-Penrose pseudo-inverse) function, which finds the least squares solution.

### Constrained numerical optimization: .optim_solve()

Linear algebra cannot provide a direct solution in the case where nonlinearities exist in the forward model. An example of this occurs when the spectrum of the primaries changes as a function of their intensity. In this case, it is still possible to obtain a solution through a numerical search procedure by solving a multi-dimensional error minimization problem of the form:

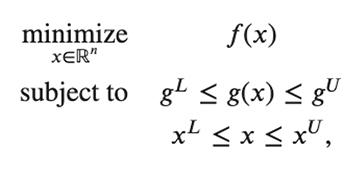

*Equation 2*. Constrained numerical optimization.
where *x* ∈ *R^n^* are the optimization variables (i.e., the primary input weights) whose lower and upper bounds, *x^L^* and *x^U^*, are between 0 and 1 to ensure that the solution is within the gamut of the device; *f*(*x*) is the objective function (which in this case is a class method that is conditioned through setting the properties outlined above) that aims to maximize contrast on the target photoreceptor(s); and g(*x*) is a constraint function (also a class method) that calculates contrast for the nominally silenced photoreceptor(s), where *g^L^* and *g^U^* should be zero. In all cases, *x* is a vector containing the primary input weights for the proposed modulation spectrum (and the background spectrum if it is not specified).

By default, *.optim_solve()* performs a local optimization from a random starting point on the function landscape using *scipy*’s (Virtanen et al., 2020) *minimize* function with the *SLSQP* (Sequential Least Squares Quadratic Programming) solver. The method may also be instructed via a keyword argument to perform a global search with *scipy*’s *basinhopping* algorithm, in which case a series of local optimisations are performed in conjunction with a global stepping algorithm and the best solution returned (Olson et al., 2012). Optimization parameters are preconfigured with sensible values, but may be changed via keyword arguments, and for full flexibility one can set up an optimization externally and pass it the objective and constraint functions from the problem class. As well as permitting the solution to silent substitution problems in cases where there are nonlinearities in the forward model, *.optim_solve()* is also a convenient way to identify background and modulation spectra that maximize unipolar amplitudes along a particular direction in photoreceptor space.

## Discussion

Silent substitution is a powerful technique for stimulating individual classes of retinal photoreceptor (Estévez & Spekreijse, 1982), but theoretical confounds and technical challenges often stand in the way of its effective application (Spitschan & Woelders, 2018). In this Methods paper we have described *PySilSub*, a new and open-source Python toolbox for performing the method of silent substitution in vision and nonvisual photoreception research. The core features described—*ColorimetricObserver*, *StimulationDevice*, *SilentSubstitutionProblem*—treat with separate problem domains of the method of silent substitution and ultimately work together to relieve the computational burden and make it easy to define, solve and visualize observer- and device-specific problems. Whilst there is plenty of scope for extensions and improvements with respect to efficiency and organization, the names and primary functions of these core features will not change in future updates to the toolbox.

The *ColorimetricObserver* code computes and stores photoreceptor action spectra appropriate to observer age (20 to 80 years) and stimulus field size (1° to 10°) in compliance with the CIEPO06 (CIE, 2006) and CIE S 026 (CIE, 2018) observer models. Previous studies vary considerably in the action spectra that are assumed for observers (Spitschan & Woelders, 2018, Table 1), which may in part be due to the difficulty in obtaining full CIE-compliant sets. Although versions of these data are maintained at various locations on the web (e.g., www.cvrl.com) or can be accessed or computed from openly available software implementations (Asano et al., 2016; Price et al., 2020; Smet, 2020), we could not find a single implementation providing a full set of photoreceptor action spectra—cones, rods and ipRGCs—appropriate to a given age and field size. Developing our own implementation afforded native support for the vast majority of human colorimetric observers. To our knowledge, this is one of few open-source methods of obtaining photoreceptor action spectra appropriate to observer age and stimulus field size and therefore has broad relevance in color and vision science, not just silent substitution. Future improvements to the *ColorimetricObserver* code may involve increasing flexibility of the wavelength sampling specification for computed action spectra, new methods for applying filter functions, and support for conversions between the radiometric and photon systems.

The main requirement for using the toolbox in one’s own research is a set of accurate calibration spectra for a multiprimary stimulation system. With these spectra, the *StimulationDevice* code can act as a forward model for the device. Various methods allow for output to be predicted for both independent primaries and multiple primaries. By default, α-opic irradiances are computed for the CIE Standard Physiological Observer, but if a custom observer is assigned, the appropriate action spectra will be used for the calculations. Overall, this part of the toolbox is best thought of as a predictive engine that performs the basic computations underlying silent substitution. As such, the output is only as good as the input, and it is important to ensure that the calibration spectra are representative of what an observer actually sees, that they are obtained with a wavelength- and irradiance-calibrated spectrometer, and that the wavelength sampling resolution agrees with that of the observer action spectra. Currently there is limited support for designing and optimizing background spectra for specifying background luminance and chromaticity, but existing functionality could be extended in future updates, or the matter outsourced to packages that already specialize in this area (e.g., LuxPy: Smet, 2020).

*SilentSubstitutionProblem* brings everything together and serves as an interface for defining, solving and visualizing silent substitution problems for a given observer and stimulation device. A problem is first conditioned by specifying which photoreceptors to ignore, silence and target, and optionally a background spectrum, the target contrast, and any restrictive bounds on the primary inputs. A solution to the problem is then obtained via a call to one of two solver methods based either on linear algebra or on constrained numerical optimization. It is important to note that the behavior of the solvers depends on the values that were set for the aforementioned properties. This is an intuitive, albeit non-standard, design pattern that facilitates quick and easy experimentation as compared to calling functions with long lists of arguments. The predicted spectral power distribution, chromaticity coordinates, and α-opic irradiances for the background and modulation spectra of a solution can be visualized via a single method call (e.g., see Figure 7) and likewise converted to native device settings when ready for use in an experiment.

The ability to ignore photoreceptors has practical advantages. For example, if studying cone-mediated vision with a standard RGB display, one could ignore both rods and melanopsin to precompute cone contrast settings for use with stimulus display software such as the Psychophysics Toolbox (Brainard, 1997; Kleiner et al, 2007) or PsychoPy (Peirce et al, 2009). If performing silent substitution in dichromats (e.g., Huchzermeyer et al., 2018; Kremers et al., 1999; Usui et al., 1998) one can simply ignore the non-functioning cone class. Similarly, if administering stimuli in the photopic range, one may wish to ignore rod photoreceptors, which are often assumed to be incapable of signaling above certain light levels (see Adelson, 1982; Aguilar & Stiles, 1954). In latter cases, ignoring the inactive photoreceptor(s) will allow for more contrast to be directed towards the targeted photoreceptor(s). Imposing limits on the primary bounds is another convenient feature that can be used to ensure that solutions avoid the hard edges of a device’s gamut, which may recede as the device ages (this is especially the case with LED-based systems).

The linear algebra and optimization solvers are complementary, but each comes with drawbacks and scope for improvement. Although solving algebraically is fast and replicable, it currently requires a background spectrum to be specified and is not suited to finding the maximum contrast available in the gamut of the device. The algebra of the toolbox could be improved in future updates by integrating more elegant examples from the literature (e.g., Evéquoz et al., 2021; Nugent et al., 2023) or by embracing the “A-matrix” approach (see Cao et al., 2015; Nugent et al., 2023) for individual observers and at a more fundamental level. Optimization on the other hand is good for finding maximum contrast but can be slow (especially with more primaries) and may give different solutions each time if the initial starting point is not fixed. Improving the speed of optimizations and supporting additional constraints (e.g., background luminance, chromaticity) may prove fruitful directions for future developments.

As a final note, we stress that *PySilSub* is a generic toolbox for dealing with the computational aspect of a challenging vision science technique. Researchers should take the usual care when designing, planning, and interpreting the results of an experiment. Accurate calibration spectra are an essential requirement to valid solutions, and it is always prudent to obtain validation measurements for a solution to ensure that the actual α-opic irradiances are as predicted by the toolbox. When interpreting the results of an experiment, researchers should acknowledge that silencing photoreceptors at the pigment level does not necessarily silence photoreceptor output (Kamar et al., 2019). The complex circuitry of the retina is such that cone cells receive feedback from other cells via horizontal cells, which means that receptor output is not dependent on quantum catch alone. That said, there is converging evidence from human experiments that the method of silent substitution can yield highly specific, photoreceptor-level characterizations. In their recent protocol paper, Nugent et al. (2023) provide a comprehensive treatment of the method of silent substitution covering also display development, calibration, and potential methods to fine-tune stimuli to individual-observer characteristics.

To conclude, we have developed an open-source Python toolbox for implementing the method of silent substitution in vision and nonvisual photoreception research. For further information on the toolbox, we refer readers to its GitHub repository (https://github.com/PySilentSubstitution/pysilsub) and online documentation pages (https://pypi.org/project/pysilsub/). Feedback and contributions in the form of issues and pull requests are warmly encouraged.

## Acknowledgements

The work was supported by funding from the Biotechnology and Biological Sciences Research Council (BBSRC: BB/V007580/1), the Engineering and Physical Sciences Research Council (EPSRC: EP/S021507/1), the Welcome Trust (204686/Z/16/Z) and John Fell OUP Research Fund, University of Oxford (0005460).

